# *metaboprep*: an R package for pre-analysis data description and processing

**DOI:** 10.1101/2021.07.07.451488

**Authors:** David A Hughes, Kurt Taylor, Nancy McBride, Matthew A Lee, Dan Mason, Deborah A Lawlor, Nicholas J Timpson, Laura J Corbin

**Affiliations:** MRC Integrative Epidemiology Unit at the University of Bristol, UK; Population Health Science, Bristol Medical School, University of Bristol, UK; NIHR Bristol Biomedical Research Centre, University of Bristol, Bristol, UK; Bradford Institute for Health Research, Bradford Teaching Hospitals NHS Foundation Trust, Bradford BD9 6RJ, UK

**Keywords:** Metabolomics, mass spectrometry, nuclear magnetic resonance, evolutionary biology, epidemiology, cohort, pipeline, ALSPAC, BiB

## Abstract

**Motivation:** Metabolomics is an increasingly common part of health research and there is need for pre-analytical data processing. Researchers typically need to characterize the data and to exclude errors within the context of the intended analysis. While some pre-processing steps are common, there is currently a lack of standardization and reporting transparency for these procedures.

**Results:** Here we introduce *metaboprep*, a standardized data processing workflow to extract and characterize high quality metabolomics data sets. The package extracts data from pre-formed worksheets, provides summary statistics and enables the user to select samples and metabolites for their analysis based on a set of quality metrics. A report summarizing quality metrics and the influence of available batch variables on the data is generated for the purpose of open disclosure. Where possible, we provide users flexibility in defining their own selection thresholds.

**Availability and implementation:** *metaboprep* is an open-source R package available at https://github.com/MRCIEU/metaboprep

## 1 Introduction

In the last decade, the study of chemical products arising from biological processes has moved from chemometrics to epidemiology (Ala-Korpela, 2015). In particular, the use of metabolomics as a functional read-out of an individual’s current health is becoming increasingly popular (Miggiels *et al*., 2019). With rapid advances in technology and bioinformatics enabling the quantification of hundreds or even thousands of metabolites from a single biological sample, there is potential for these measurements to reveal valuable insights into biology and health. Both mass spectrometry (MS) and nuclear magnetic resonance (NMR) are common technologies used in the metabolomics field. Typically, laboratories have their own established protocols in sample preparation, generation of standards and controls, and corrections for instrument and run day variability. As a result, researchers are now able to access high quality curated metabolomics data at scale.

Subsequent to the data generation by core facilities and prior to statistical analysis, researchers perform a series of data characterization and pre-analytical preparation steps. These may include (1) the identification of samples of poor quality, (2) the identification of metabolites that have unfavourable statistical properties and/or may not provide sufficient data for study analyses, and (3) to characterize statistical properties of the data that may be relevant to downstream analyses. The latter is needed to help inform decisions involving data normalizations, transformations and analytical considerations that revolve around missing data.

Based on both our own experience and emerging literature published in this area (Barnes, 2020; Monnerie *et al*., 2020), it is clear that approaches for post-acquisition, pre-analytical data processing are varied both within and across analytical platforms. The general lack of methodological standardization makes combining and comparing data and results across studies difficult, thus impairing cross-study inference. Into this context, papers and researchers have recently called for standardization and transparency in reporting of metabolomic studies (van Roekel *et al*., 2019; Long *et al*., 2020; Karaman, 2017; Begou *et al*., 2018; Playdon *et al*., 2019). To some extent, this situation mirrors that seen in the field of genomics a decade ago; here, researchers responded with the development of standard protocols supported by open-source software tools such as EasyQC (Winkler *et al*., 2014). Under the assumption that, as in genomics, collaboration and independent replication will be key in the utilization of metabolomics data going forward, it is important that this field progresses in optimizing workflows and recognizing where consistency can be achieved. In addition, efforts need to be made to improve transparency in the reporting of pre-analytical data processing.

This paper introduces *metaboprep*, an R package developed to help those working with curated metabolomics data to achieve transparent and informed processing of their study sample data prior to statistical analysis. The package provides a detailed summary of the data, highlighting properties relevant both to setting sample/metabolite filtering criteria and to downstream analytical choices. *metaboprep* can process any flat text data file containing curated metabolomics data with minimal formatting. In addition, the *metaboprep* package is currently able to process data as supplied by two of the main biotech companies operating in this sector – ^1^H-NMR data from Nightingale Health© (Helsinki, Finland) and ultra-high-performance liquid chromatography-tandem mass spectrometry (UPLC-MS/MS) data from Metabolon (Research Triangle Park, NC, USA). We demonstrate the use of *metaboprep* using the Born in Bradford (BiB) cohort, including 1,000 pregnant women with UPLC-MS/MS data (Metabolon), and The Avon Longitudinal Study of Parents and Children (ALSPAC), a birth cohort with 3,361 samples collected during early adulthood and analysed by NMR (Nightingale Health).

## 2 Materials and methods

### Overview

*Metaboprep* is an R package designed to standardise the steps involved in preparing population level metabolomics data sets for statistical analysis. It was written using R (version 3.6.0) (R Core Team, 2019), is dependent upon R version 3.4.0 or greater, and is available on GitHub (https://github.com/MRCIEU/metaboprep). A README is available on the *metaboprep* GitHub page that provides detailed instructions for running the *metaboprep* pipeline and functions used within. All analyses performed in this manuscript used R (version ≥ 3.4.0); code is available on the repository providing a walk-through of the *metaboprep* package. Data used here is available upon application from BiB and ALSPAC, and an example data set is also available on the GitHub repository.

### Data processing and filtering pipeline

When run in its entirety, the *metaboprep* package performs five main actions: (1) extracts and processes (un)targeted metabolite data from source files, saving datasets in a standard tab-delimited format for use elsewhere; (2) generates summary statistics from the initial raw data set which are exported to a standard tab-delimited text file; (3) performs sample and metabolite filtering according to user-defined thresholds and using a standard pipeline; (4) repeats the generation of summary statistics but on the filtered data set; (5) and finally summarises the data in a PDF report while also reporting on the influence of available batch variables.

An overview of the workflow is shown in **Figure 1** and a brief description given below. A log file is generated detailing each step taken in the pipeline including the filtering thresholds defined by the user and the number of samples or metabolites excluded at each step. We do not normalize or transform the data in any way for the user, nor do we perform any imputation. This is because we feel decisions around whether and how to do these depend on specific analyses for specific research questions and the aim of *metaboprep* is to undertake processes that we consider valuable to be completed on metabolomics data before any main analyses. We do report on the normality (Shapiro-Wilk W-statistic) of log-transformed data in the PDF report to help inform the user on its appropriateness in parametric analyses.

**Figure 1:**
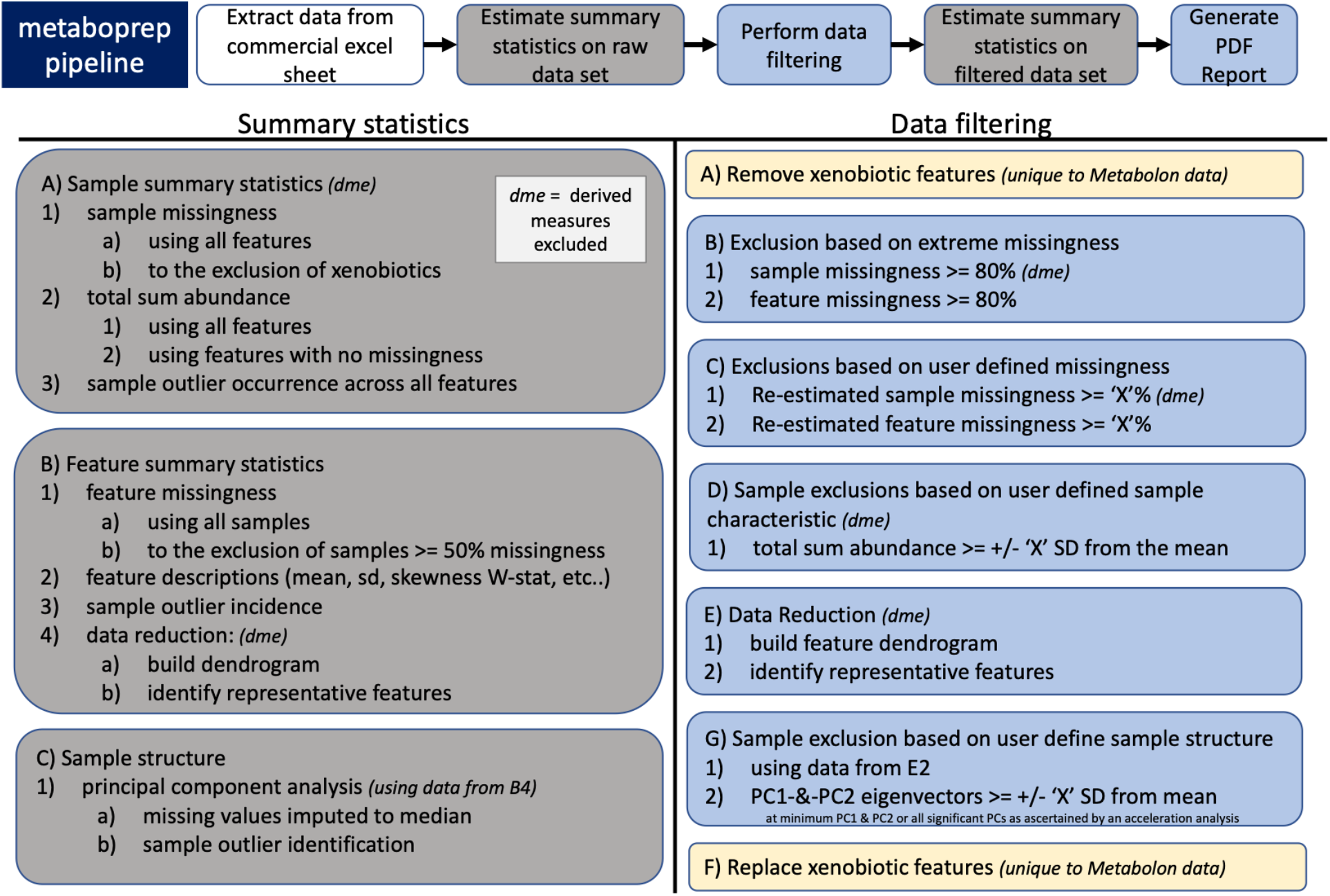
Workflow. Brief description of the *metaboprep* pipeline. Along the top, the five primary steps the pipeline takes are outlined. The column on left provides an outline of the steps for the generation of summary statistics while the right provides an outline of the steps taken for sample and metabolite filtering. Common abbreviations used are: ‘dme’ for derived measures excluded; SD for standard deviations; ‘X’ which denotes a threshold variable that is defined by the user in the pipeline parameter file; PC for principal components.

### Running the R package

The package can be run via the command line using a parameter file but can also be run in an interactive mode using the functions built within. The parameter file contains key information for running the pipeline including the project name, the path to the source data directory, the input file names, the platform the metabolomics data is derived from and the preferred filtering thresholds. Thresholds to be provided are: (1) the fraction of metabolite missingness retained, (2) the fraction of sample missingness retained, (3) the total sum abundance (TSA) threshold in standard deviations from the mean, (4) the principal component (PC) threshold for samples in standard deviations from the mean, and (5) if derived variables should be excluded (TRUE or FALSE).

### Data Extraction

Input data files can be in one of two possible formats, (1) excel spreadsheets as supplied by Nightingale Health or Metabolon; (2) formatted tab-delimited text files. When an excel spreadsheet is provided as the source data, the package extracts (a) the (semi-) quantified metabolite data, (b) the associated sample metadata (e.g. technical batch information, sample identifiers) and (c) the associated metabolite metadata (e.g. metabolite class/pathway, HMDB identifier) and writes each ‘raw’ (i.e. unaltered) data set to its own tab-delimited text file. Alternatively, the user can provide their (a) metabolite data, (b) optional sample metadata, and (c) optional metabolite metadata, as text files (.csv or .txt). The text files can contain metabolite abundance data from any platform.

Special considerations are made for certain metabolites when the data is derived from the commercial platforms of Metabolon or Nightingale Health. In the case of Metabolon data, metabolites labelled as “xenobiotics” are excluded from calculations relating to sample missingness and do not follow the same processing pipeline as other metabolites. Xenobiotics are treated in this way because of their typically high level of missingness (or absence), which is both expected and biologically relevant given their (predominantly) exogenous origins. In the case of Nightingale Health data, many of the data summary and filtering steps are carried out having (optionally) excluded derived measures, which include a number of measures expressed as percentages or ratios. In brief, including these measurements could, for example, artificially inflate missingness estimates and lead to unwanted exclusions. See **Discussion** for further explanation on this point.

### Data summary

A data summary is generated twice by the package. Once on the raw, unaltered dataset and again on the filtered (analysis ready) dataset. The summary includes a series of sample- and metabolite-based summary statistics. We use ‘sample’ here as a generic term that in many studies will mean the same as participant as analyses to generate metabolite data will have been run on one sample per participant. However, as some studies will have repeat samples drawn over time from the same participants, sample, as used here means each individual sample on which metabolites are measures. Sample-based summary statistics include (1) an estimate of sample missingness, calculated as the proportion of missing (‘NA’) data and (2) total sum abundance (TSA; often referred to as total peak area (TPA) for those familiar with mass spectroscopy data), calculated for each sample by summing values across all metabolites. TSA provides an estimate of the total (measured) metabolite concentration in the sample. Sample missingness is estimated using (a) all metabolites and again (b) to the exclusion of any defined list of metabolites, such as xenobiotics or derived measures. Sample TSA is estimated using (a) all metabolites and (b) again using only those metabolites with no missing data. Additionally, an outlier occurrence count, or an integer count of the total number of times an individual’s metabolite value is more than five standard deviations from the mean metabolite concentration, is calculated and provided in the sample-based summary statistics file. Finally, subsequent to the estimation of metabolite summary statistics and the identification of representative metabolites, sample PC’s are estimated, and the top 10 PCs provided in the summary data. This latter step is detailed below. Metabolite-based summary statistics include metabolite missingness, sample size (n) or the count of individuals without missing (‘NA’) data, and numerous other descriptive statistics including mean, standard deviation, skew, and the coefficient of variance. A direct measure of each metabolites’ data distribution conformity to normality is provided by an estimate of Shapiro’s W-statistic, provided for both untransformed and log10 transformed data distributions. In addition, for each metabolite a count of the number of outlying samples is provided as a further indication of skewness.

For the purposes of defining correlation structure and identifying a subset of approximately independent or ‘representative’ metabolites from the complete set of metabolites, a data reduction step is performed. These analyses provide users with a count of the effective number of metabolites in their data set, which could be used for multiple testing correction, as well as a list of ‘representative’ metabolites. In the case of Nightingale Health NMR data or other data containing ratios of metabolites, derived measures can be excluded from this step. Further, data are restricted to common metabolites such that only those that are (a) variable and (b) have less than or equal to 20% missingness are included. A dendrogram is then constructed (‘stats’ package hclust() function, with method ‘complete’) based on a Spearman’s rho distance matrix (1-|Spearman’s rho|). A set of ‘k’ clusters (groups of similar metabolites) are identified based on a user-defined tree cut height (default 0.5 and equivalent to a Spearman’s rho of 0.5), using the function cutree() from the ‘stats’ package. For each ‘k’ cluster the metabolite with the least missingness is then tagged as the representative metabolite for that cluster. Representative metabolites are identified by 1’s in the metabolite summary statistics file in the column “independent_features_binary”.

A principal component analysis (PCA) is conducted to evaluate inter-individual variability in metabolomic profiles. This sample-based analysis uses only the subset of representative metabolites previously identified. Strictly for the purposes of deriving the PCs, missing values are imputed to the median and data then standardised (z-transformed) so that the mean equals zero and the standard deviation equals one for each metabolite. The variance explained for each PC is extracted and an estimate of the number of PCs (*n*) to retain is estimated, by both the acceleration factor and parallel analysis with the function nScree() from the ‘nFactors’ R package. The estimate of *n* derived by the acceleration factor, with a defined minimum of two, is used to identify sample outliers or those that deviate too far from the mean on those *n* PCs. By default, the outlier threshold is defined as five standard deviations from the mean, but this can be set by the user.

The summary statistics described above are written to two tab-delimited text files, one for samples and one for metabolites, and additionally once for the raw dataset and once for the filtered dataset. In addition, key statistics are reported (including graphically) in a PDF report (see below).

### Data filtering

The next step in the pipeline is to derive a version of the metabolomics dataset which has undergone sample and metabolite filtering according to the user specifications provided in the parameter file. The first step is to remove, first samples, and then metabolites with extremely high rates of missingness (>=80%). In the second step, missingness is recalculated and sample and metabolite exclusions are made according to the user-defined thresholds for missingness with a default suggested value of 0.2 or 20%. Sample exclusions are then made on TSA, using only metabolites with complete data, according to user-defined thresholds with a suggested default of five standard deviations from the mean. Then, using all remaining samples and metabolites, the clustering and sample PCA steps described previously (as part of the data summary) are repeated. Results of the PCA are used to identify sample outliers for exclusion, defined as those that lie more than the user-defined threshold from the mean on *n* PCs where *n* is defined by the acceleration factor (as described previously). The default suggested threshold value is five standard deviations from the mean. This post-filtering version of the metabolite data is then passed back through the data summary procedure described above and finally exported in flat text format.

The most appropriate thresholds for sample and feature missingness will depend on both the total sample size and the intended analysis and we recommend users consider carefully the thresholds they set. Additionally, sample metabolome profile exclusions (TSA and PCA) are set at five standard deviations from the mean, as we have observed this to be a reasonable threshold to exclude samples that perform poorly, when sampling a random presumptively healthy population. If sampling something similar to a case-control study design where extremes are perhaps expected or indeed a study sample with known substructure (e.g. different sample types) it would be advisable to evaluate the distributions in the packages PDF report (see below) and consider modifying these parameters.

### PDF report

The standardised PDF report (designed for inclusion in papers in order to facilitate data description and hence transparency) includes the project name, the platform, a workflow image, contact information for feedback, data summaries, and analysis of batch effects on key properties of the data – missingness and TSA. The data summary includes (1) an overview of the raw dataset: (1a) a visual of missingness in the data matrix, (1b) samples and metabolite missingness distributions; (2) an overview of the filtering steps: (2a) an exclusion summary, (2b) metabolite data reduction summary, and (2c) a PC plot illustrating sample structure and identifying potential sample outliers; (3) a summary of the filtered dataset: (3a) count of remaining samples and metabolites, (3b) distributions for sample missingness, metabolite missingness and TSA, (3c) a metabolite clustering dendrogram highlighting the representative metabolites, (3d) a metabolite data reduction summary, (3e) a PC plot of sample structure, (3f) histograms for Shapiro W-statistic estimates across untransformed and log10 transformed metabolite abundances, and (3g) a summary of sample and metabolite outliers. The report also includes an evaluation of the relationship between key sample properties (missingness rates and TSA) and potential batch variables, as provided by the user. Such variables include sample storage box identifier, run day, super- and sub-pathway, sampling data and time, and MS run mode. How these batch variables associate with missingness and TSA is illustrated in a series of boxplots that include an estimate of the variance explained by the batch, derived from a univariate analysis of variance and estimation of eta-squared using sums of squares. In addition, all the identified batch variables are placed in a type II multivariate analysis of variance and again the variance explained by each is summarised by the eta-squared statistic.

A power analysis for continuous and binary traits is provided based on the sample size of the dataset and using functions from the ‘pwr’ R package. If researchers are interested in the relationship between metabolites and a continuous trait (for example, weight), power estimates are provided assuming a general linear model, whereas for the case of binary analyses (e.g. case/control) calculations are based on a two-sample t-test (allowing unequal sample sizes). The aim of these power calculations is to demonstrate the loss of power that can be expected as a result of varying degrees of missing data (i.e. as actual sample size decreases). During the generation of the main PDF report, a second PDF is written that contains, for each metabolite, a scatter plot identifying outlying data points and a histogram that includes selected summary statistics. Together, these two PDFs provide a quick overview and reference for the dataset.

### Example datasets

#### *Born in Bradford* – Mass Spectrometry

The Born in Bradford (BiB; https://borninbradford.nhs.uk/) study is a population-based prospective birth cohort based in Bradford, United Kingdom. Full details of study methodology have been reported previously (Wright *et al*., 2013). Ethical approval for the study was granted by the Bradford National Health Service Research Ethics Committee (ref 06/Q1202/48), and all participants gave written informed consent. For the data used in this example, women of White British (N=500) or Pakistani (N=500) ancestry were selected to have samples analysed on the basis of their having complete data on a set of pre-specified variables, including valid pregnancy fasting and post-load glucose measures, and both them and their index child having genome-wide data available (as described previously (Taylor *et al*., 2020) and in **Supplementary Figure S1**). Samples were collected during pregnancy at around 24-28 weeks’ gestation. Participant characteristics are shown in **Supplementary Table S1**.

The untargeted metabolomics analysis of over 1,000 metabolites was performed on these samples at Metabolon, Inc. (Durham, North Carolina, USA) on a platform consisting of four independent ultra-high-performance liquid chromatography-tandem mass spectrometry (UPLC-MS/MS) runs. Detailed descriptions of the platform can be found in **Supplementary Methods** and in published work (DeHaven *et al*., 2010; Evans *et al*., 2009; Taylor *et al*., 2020). The resulting datasets comprised a total of 1,369 metabolites, of which 1,000 were of known identity (named biochemicals) at the time of analysis. This dataset will be referred to throughout as BiB_MS-1.

#### Avon Longitudinal Study of Parents and Children - NMR

The Avon Longitudinal Study of Parents and Children (ALSPAC; http://www.bristol.ac.uk/alspac/) is a prospective birth cohort study, based in the former region of Avon, United Kingdom. Detailed information about the methods and procedures of ALSPAC can be found in **Supplementary Methods** and in published work (Fraser *et al*., 2013; Boyd *et al*., 2013; Northstone *et al*., 2019). Ethical approval for the study was obtained from the ALSPAC Ethics and Law Committee and the Local Research Ethics Committees (a full list of all ethical approvals relating to ALSPAC are available online: http://www.bristol.ac.uk/alspac/researchers/research-ethics/). Specifically ethical approval for the clinic in which samples were collected for this work was granted by the National Research Ethics Service Committee South West – Frenchay (14/SW/1173). Consent for biological samples has been collected in accordance with the Human Tissue Act (2004).

NMR-derived metabolomics data were derived for 3,361 EDTA-plasma/serum samples collected from 3,277 unique individuals during the age 24 years clinic visit. Participant characteristics are shown in **Supplementary Table S2**. Quantification of selected circulating lipids, fatty acids, and metabolites was performed using a 1D proton (1^H^) NMR spectroscopy-based platform from Nightingale Health (Helsinki, Finland). Spectra were acquired using standardised parameters using two NMR experiments or ‘molecular windows’ to characterise lipoproteins, low molecular weight metabolites and lipids. Further information relating to the data derivation can be found in **Supplementary Methods** and has been described previously (Inouye *et al*., 2010; Soininen *et al*., 2015, 2009). Raw metabolomics data pre-processing and quantification were as previously described (Soininen *et al*., 2015; Inouye *et al*., 2010; Soininen *et al*., 2009). The resulting dataset comprised a total of 225 metabolites (including 78 derived measures); this dataset will be referred to throughout as ALSPAC_F24.

## 3 Results

We used data from two established population-based cohorts (BiB_MS-1 and ALSPAC_F24) and two different analytical platforms (Metabolon and Nightingale Health) to demonstrate the utility of *metaboprep*. The summary PDF reports generated for each dataset can be found in **Supplementary Data.** The single core machine run times for the datasets were 3 and 10 minutes for ALSPAC_F24 and BiB_MS-1, respectively. An overview of each dataset based on the summary statistics generated by *metaboprep* prior to filtering are shown in **Table 1.** The choice of user-defined thresholds used in our analyses and the resulting exclusions made are summarised in **Table 2**. Based on the thresholds used here, 11 and 6 samples were excluded from BiB_MS-1 and ALSPAC_F24, respectively. The sample exclusions made on the basis of the PCA in BiB_MS-1 relate to an underlying sub-structure in these data made obvious by the *metaboprep* steps (**Figure 2**) (see **Discussion**). No metabolites were excluded from ALSPAC_F24 whilst metabolite missingness criteria resulted in 24% of metabolites being excluded from BiB_MS-1.

**Table 1.** Summary statistics by dataset (pre-filtering) Summary statistics for the initial, raw (pre-filtered) BiB_MS-1 and ALSPAC_F24 datasets. The table provides details on the platform, sample size, sample and metabolite missingness, total sum abundance (TSA) for samples, and outlier counts, the percent of metabolites that may be considered normal distributed and an estimate of the number of representative metabolites in the data set. *calculated after the exclusion of derived variables in the Nightingale Health dataset and of xenobiotics in the Metabolon dataset.

| Summary statistic | BiB_MS-1 | ALSPAC_F24 |
| --- | --- | --- |
| Platform | Metabolon | Nightingale Health |
| No. of samples | 1000 | 3361 |
| No. of metabolites | 1369 | 225 |
| **Sample statistics** |  |  |
| % sample missingness*<br>(min, median, max) | 11.85, 18.45, 26.81 | 0.00, 0.00, 12.24 |
| Total sum abundance at complete metabolites (min, median, max) | 4.23, 6.79, 9.91 ($\times 10^{12}$) | 1.66, 2.08, 2.46 ($\times 10^5$) |
| Count of outlying data points per sample<br>(min, median, max) | 0, 2, 81 | 0, 0, 44 |
| **Metabolite statistics** |  |  |
| % metabolite missingness (min, median, max) | 0, 2.6, 100 | 0.00, 0.00, 1.73 |
| Count of outlying data points per metabolite<br>(min, median, max) | 0, 2, 16 | 0, 2, 97 |
| % with W statistic $\geq 0.95$ | 16.08 | 42.22 |
| % whose W-statistic decreases following $\log_{10}$ transformation | 9.56 | 44.89 |
| No. of representative metabolites | 512 | 24 |

**Table 2.** Results of sample and metabolite filtering based on default exclusion thresholds. PCA, principal component analysis; SD, standard deviations. ^a^Calculated after excluding metabolites in the xenobiotic class from Metabolon data and derived measures from Nightingale Health data; ^b^derived from complete metabolites only; ^C^excluding metabolites with >20% missingness; ^d^using the representative metabolites only and excluding on the number of PCs determined by the acceleration factor with a minimum of two PCs; *user defined threshold. Rows in blue are sample filtering steps.

| Filtering step | Exclusion threshold | BiB_MS-1 | ALSPAC_F24 |
| --- | --- | --- | --- |
| Raw dataset (pre-filtering) |  | 1000 samples<br>1369 metabolites | 3361 samples<br>225 metabolites |
| 1. extreme sample missingness<sup>a</sup> | $\geq 80\%$ | 0 | 0 |
| 2. extreme metabolite missingness<sup>a</sup> | $\geq 80\%$ | 96 | 0 |
| 3. sample missingness<sup>a*</sup> | $\geq 20\%$ | 0 | 0 |
| 4. metabolite missingness<sup>a*</sup> | $\geq 20\%$ | 236 | 0 |
| 5. sample total sum abundance<sup>b*</sup> | $> 5SD$ | 0 | 2 |
| 7. PCA outliers<sup>c,d*</sup> | $> 5SD$ | 11 | 4 |
| Final dataset (post-filtering) |  | 989 samples<br>1037 metabolites | 3355 samples<br>225 metabolites |

**Figure 2:**
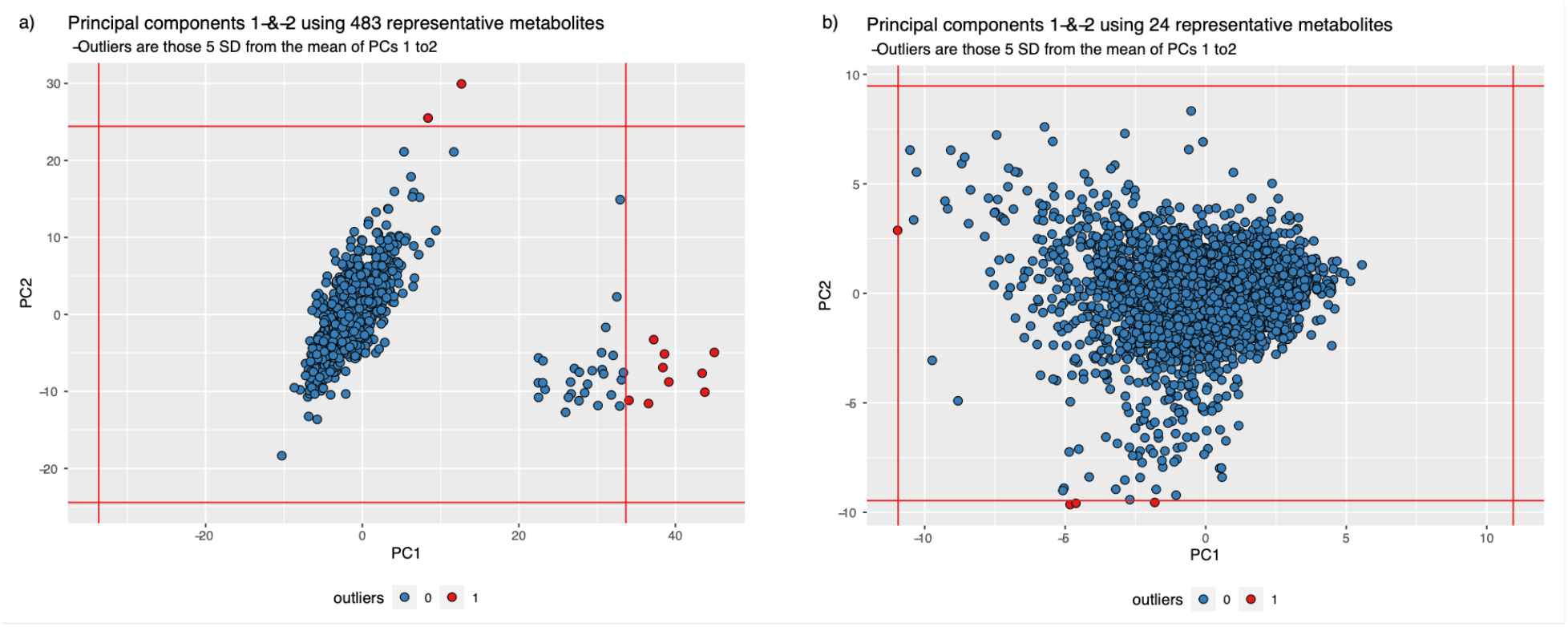
Principal component analysis to show sample structure. Principal components one and two of a) BiB_MS-1 dataset; b) ALSPAC_F24 dataset. Red vertical and horizontal lines represent five standard deviations from the mean on both PC axes, used to identify outliers in the dataset.

## 4 Discussion

In this paper, we have presented *metaboprep*, an R package for use by researchers working with curated, high quality, metabolomics data and developed in the context of population health research. The package enables metabolomics data from different platforms to be extracted, processed, summarised and prepared for subsequent statistical analysis within a standardised and reproducible workflow. This work was motivated by the need for increased consistency and transparency in the pre-analytical processing of data across cohorts and studies, but also acknowledges that a ‘one size fits all’ approach is unlikely to be feasible given the range of study designs being employed. Metabolomics is a growing field within population sciences, with application to a vast array of hypotheses. As such, research groups have differing approaches to data preparation, which can make results hard to interpret and compare across studies. It is important to understand the properties of metabolomics data in order that suitable pre-analytical processing steps can be performed, and downstream analytical results interpreted appropriately.

In the proposed pipeline, considerations were made for two specific conditions within two platforms currently available. It is difficult to mitigate against future developments, but the *metaboprep* approach is able to accommodate specific flags as they appear. In this case, xenobiotics - common in the Metabolon dataset, and derived variables - common in the Nightingale Health dataset. Xenobiotics are exogenous metabolites (i.e. not produced by the body), such as drug compounds and a quantified measure indicates presence of the exogenous compound. Consequently, they can have very high rates of missingness, while still being critically informative to a study as the majority with missing data will be a true ‘no’ for exposure to the exogenous compound. For this reason, we do not exclude xenobiotics on the basis of high missingness but would advocate affording them special consideration in any downstream statistical analyses. For example, these metabolites might best be evaluated within a presence/absence framework rather than by analysis of relative abundance. In data from Nightingale Health, derived variables are metabolite traits that are a summary of two or more other metabolites (possibly already represented in the dataset) or ratios of two or more metabolites. These variables can introduce bias in estimates of sample missingness (where a single metabolite is missing, any derived measures based on that metabolite will also be missing) and may not be appropriate to retain when identifying a set of representative metabolites for the data set. We allow the user to include or exclude the derived variables in the pipeline at their discretion.

One of the most commonly implemented pre-analytical steps is filtering based on missingness. Missingness is defined as the proportion of data with no value and can vary hugely within a dataset (0-99.9% missing). Typically, researchers filter metabolites on missingness to remove metabolites that exhibit evidence of technical error, or where the proportion of missingness introduces downstream analytical difficulties. Conversely, the filtering of samples based on missingness helps identify samples that may have been of poor quality or mishandled before or during the metabolomic assay(s). However, without external data the nature of the missingness, and the extent to which removal of samples or metabolites introduces more or less bias than not excluding these (and possibly imputing missing data) is unclear (Hughes *et al*., 2019) and will vary by sample size and the intended main research questions. Deciding upon appropriate missingness thresholds can be critical to a study and some caution and consideration is warranted. This results from the variety of reasons for missing data in this context – e.g. the technology, signal to noise ratios, signal intensity, error (Do *et al*., 2018) (for further discussion on missingness see **Supplementary Discussion**). Crucially, missingness might also represent true absence and thus be informative for some biological hypotheses, for example, differential missingness by class (e.g. case/control status or sex). For this reason, our workflow allows uses to define the thresholds they want to apply for missing samples and metabolites, the thresholds for these two can be different and either or both can be zero (no exclusions based on missing data). This allows researchers to repeat the workflow with different thresholds to explore the extent that these influence main analysis results.

TSA is a sample-based metric estimated for the purposes of identifying samples with broad quality issues, such as handling errors (i.e. differing concentrations of sample) and is calculated by summing values across all metabolites. This metric is, by definition, correlated with missingness rates, so is estimated a second time here using only complete metabolites with this latter metric being used in the exclusion step. In order to guard against selection bias, the implementation of this exclusion step should be considered carefully and within the context of the study design. There may be situations whereby a high (or low) TSA is indicative of a true biological state, rather than of any technical issue. For example, if the coverage of the metabolomics platform is skewed towards a class of metabolite, e.g. lipids, then certain characteristics of individuals in the study sample may be correlated with the TSA measure, e.g. body fat percentage. Alternatively, if a study design were to include data from various tissues, then the TSA distribution may be bimodal and basing exclusions on standard deviations from the mean may be difficult if not inappropriate. For these reasons, the TSA distribution is provided in the PDF report for assessment by the user who may then choose to explore the sensitivity of downstream analyses to the application of different thresholds.

The proposed workflow provides information relating to structure within the study sample. This is done by implementation of sample-based PCA with summary data provided in the summary statistic files and corresponding plots for visual inspection. Only metabolites with limited missingness (<20%) are included in these analyses to avoid the need to implement a probabilistic PCA whilst limiting the introduction of error by the simplified (median-based) imputation – an imputation used strictly for deriving PCs. Furthermore, data reduction to remove highly correlated metabolites is considered necessary to ensure that the estimated PCs are not driven by any common, highly correlated metabolite classes, pathways, or clusters. Taking this approach, the PCs should provide an equally represented, broad perspective of variation in the data. If a sample is mishandled, the assumption would be that all of the assayed variables would be perturbed, and this would be evident in the PCA. Just as discussed with missingness and TSA metrics, proper consideration for thresholds is important here too. Outliers may be biologically relevant and gross structure may be present if multiple tissues, populations, or species are sampled. If that is the case, just as for TSA, then thresholding on standard deviations from the mean may be a difficult if not inappropriate filtering step. However, if you are anticipating a homogenous sample but observe clustering (as in **Figure 2**), then you should attempt to identify the source of the clustering and potentially re-consider your PCA sample filtering threshold.

Three pre-processing steps that the *metaboprep* pipeline does not, currently, incorporate is modification of outlying data points (winsorization or truncation), data transformation and imputation. Each of these topics bring with them their own particular issues and considerations that are beyond the scope of the current package. We will however note that while log transformations appear to be commonly applied to metabolomics data sets - 64% of COMETS (The COnsortium of METabolomics Studies) responding cohorts claim to routinely log transform their data (Playdon *et al*., 2019) – we routinely observe that this does not always generate an approximate normal distribution and at times can make data distributions less normal. Shapiro W-statistics (a metric for normality) are provided alongside outlier flags in the summary statistic file for metabolites and the distribution of W-statistics for the raw and log-transformed data is provided in the PDF report. We encourage use of this information to aid decisions regarding the most appropriate data transformation(s), given the intended statistical analyses. These considerations should also include the research question, including whether the metabolites are exposures or outcomes, and the planned main analyses.

To date, the *metaboprep* package has only been used to process metabolomics data derived from serum or plasma, not other biological samples (e.g. urine, tissue). Whilst we do not anticipate any issues in processing data derived from other sources, users should consider carefully whether the assumptions we make are appropriate in these scenarios. The same is true if the package is used for processing small samples (n<20), where steps such as the identification of independent metabolites may not perform optimally. We reiterate that the workflow presented here does have its compromises. As highlighted above, data preparation does not end with the running of this workflow but with the careful evaluation of the data reports provided by it.

In conclusion, in the interests of open science and to encourage collaboration we present a first release of *metaboprep*, an R package that we hope to develop further in response to feedback from the community. In this paper, we have avoided making definitive recommendations regarding thresholds that should be used since these should be chosen in the context of the specific study design and research question. We encourage those working with curated metabolomics data to use our package to enhance their understanding of the characteristics of their metabolomics data, its structure and how these properties could impact on downstream statistical analyses and importantly, to report their findings alongside the results of their main analyses.

## 5 Acknowledgements

We are grateful to everyone involved in the Born in Bradford study. This includes the families who kindly participated, as well as the practitioners and researchers all of whom made Born in Bradford happen.

We are extremely grateful to all the families who took part in the ALSPAC study, the midwives for their help in recruiting them, and the whole ALSPAC team, which includes interviewers, computer and laboratory technicians, clerical workers, research scientists, volunteers, managers, receptionists and nurses. Sample processing and NMR analysis were carried out at the Bristol Bioresource Laboratory and the NMR Metabolomics facility at the University of Bristol.

## 8 List of supplementary data

**Supplementary Data 1:**

Supplementary Methods
Supplementary Discussion
Supplementary Figures
Supplementary Tables
Supplementary References

**Supplementary Data 2:** PDF report for BiB_MS-1

**Supplementary Data 3:** PDF report for ALSPAC_F24

