## Supplementary_Data_1 for "*metaboprep*: an R package for pre-analysis data description and processing"

***metaboprep:*** **an R package for pre-analysis data description and processing.**

#### Table of Contents


***Supplementary Methods*** **2**

**Metabolite data acquisition** **2**

Metabolon 2

Nightingale Health 4

**Born in Bradford** **5**

**Avon Longitudinal Study of Parents and Children** **5**

Description of study numbers 5

Data management 6

Details of ethics approvals 6

***Supplementary Discussion*** **8**

**Sources and properties of missingness** **8**

***Supplementary Figures*** **9**

**Figure S1. Born in Bradford sample selection flow chart** **9**

***Supplementary Tables*** **10**

**Table S1. Born in Bradford (BiB) dataset: Participant characteristics** **10**

**Table S2. Avon Longitudinal Study of Parents and Children (ALSPAC) dataset: Participant characteristics** **11**

***Supplementary References*** **12**

### Supplementary Methods

#### Metabolite data acquisition

##### Metabolon

The details below are as supplied by Metabolon and as described previously (Evans *et al.*, 2009; DeHaven *et al.*, 2010).

*Sample preparation:* Following receipt, samples were inventoried and immediately stored at -80$^{o}$C. All samples were maintained at -80$^{o}$C until processed. Samples were prepared using the automated MicroLab STAR® system from Hamilton Company. Several recovery standards were added prior to the first step in the extraction process for quality control (QC) purposes. To remove protein, dissociate small molecules bound to protein or trapped in the precipitated protein matrix, and to recover chemically diverse metabolites, proteins were precipitated with methanol under vigorous shaking for 2 min (Glen Mills GenoGrinder 2000) followed by centrifugation. The resulting extract was divided into five fractions: two for analysis by two separate reverse phase (RP)/UPLC-MS/MS methods with positive ion mode electrospray ionization (ESI), one for analysis by RP/UPLC-MS/MS with negative ion mode ESI, one for analysis by HILIC/UPLC-MS/MS with negative ion mode ESI, and one sample was reserved for backup. Samples were placed briefly on a TurboVap® (Zymark) to remove the organic solvent. The sample extracts were stored overnight under nitrogen before preparation for analysis.

*Quality assurance (QA)/ Quality control (QC):* Several types of controls were analyzed in concert with the experimental samples: a pooled matrix sample generated by taking a small volume of each experimental sample (or alternatively, use of a pool of well-characterized human plasma) served as a technical replicate throughout the data set; extracted water samples served as process blanks; and a cocktail of QC standards that were carefully chosen not to interfere with the measurement of endogenous compounds were spiked into every analyzed sample, allowing instrument performance monitoring and aiding chromatographic alignment. Instrument variability was determined by calculating the median relative standard deviation (RSD) for the standards that were added to each sample prior to injection into the mass spectrometers. Overall process variability was determined by calculating the median RSD for all endogenous metabolites (i.e., non-instrument standards) present in 100% of the pooled matrix samples. Experimental samples were randomized across the platform run with QC samples spaced evenly among the injections.

*Ultrahigh Performance Liquid Chromatography-Tandem Mass Spectroscopy (UPLC-MS/MS):* All methods utilized a Waters ACQUITY ultra-performance liquid chromatography (UPLC) and a Thermo Scientific Q-Exactive high resolution/accurate mass spectrometer interfaced with a heated electrospray ionization (HESI-II) source and Orbitrap mass analyzer operated at 35,000 mass resolution. The sample extract was dried then reconstituted in solvents compatible to each of the four methods. Each reconstitution solvent contained a series of standards at fixed concentrations to ensure injection and chromatographic consistency. One aliquot was analyzed using acidic positive ion conditions, chromatographically optimized for more hydrophilic compounds. In this method, the extract was gradient eluted from a C18 column (Waters UPLC BEH C18-2.1x100 mm, 1.7 µm) using water and methanol, containing 0.05% perfluoropentanoic acid (PFPA) and 0.1% formic acid (FA). Another aliquot was also analyzed using acidic positive ion conditions; however, it was chromatographically optimized for more hydrophobic compounds. In this method, the extract was gradient eluted from the same afore mentioned C18 column using methanol, acetonitrile, water, 0.05% PFPA and 0.01% FA and was operated at an overall higher organic content. Another aliquot was analyzed using basic negative ion optimized conditions using a separate dedicated C18 column. The basic extracts were gradient eluted from the column using methanol and water, however with 6.5mM Ammonium Bicarbonate at pH 8. The fourth aliquot was analyzed via negative ionization following elution from a HILIC column (Waters UPLC BEH Amide 2.1x150 mm, 1.7 µm) using a gradient consisting of water and acetonitrile with 10mM Ammonium Formate, pH 10.8. The MS analysis alternated between MS and data-dependent $MS^n$ scans using dynamic exclusion. The scan range varied slighted between methods but covered 70-1000 m/z. Raw data files are archived and extracted as described below.

*Data extraction and compound identification:* Raw data was extracted, peak-identified and QC processed using Metabolon's hardware and software. These systems are built on a web-service platform utilizing Microsoft's .NET technologies, which run on high-performance application servers and fiber-channel storage arrays in clusters to provide active failover and load-balancing. Compounds were identified by comparison to library entries of purified standards or recurrent unknown entities. Metabolon maintains a library based on authenticated standards that contains the retention time/index (RI), mass to charge ratio (*m/z)*, and chromatographic data (including MS/MS spectral data) on all molecules present in the library. Furthermore, biochemical identifications are based on three criteria: retention index within a narrow RI window of the proposed identification, accurate mass match to the library +/- 10 ppm, and the MS/MS forward and reverse scores between the experimental data and authentic standards. The MS/MS scores are based on a comparison of the ions present in the experimental spectrum to the ions present in the library spectrum. While there may be similarities between these molecules based on one of these factors, the use of all three data points can be utilized to distinguish and differentiate biochemicals. More than 3300 commercially available purified standard compounds have been acquired and registered into LIMS for analysis on all platforms for determination of their analytical characteristics. Additional mass spectral entries have been created for structurally unnamed biochemicals, which have been identified by virtue of their recurrent nature (both chromatographic and mass spectral). These compounds have the potential to be identified by future acquisition of a matching purified standard or by classical structural analysis.

*Curation:* A variety of curation procedures were carried out to ensure that a high-quality data set was made available for statistical analysis and data interpretation. The QC and curation processes were designed to ensure accurate and consistent identification of true chemical entities, and to remove those representing system artifacts, mis-assignments, and background noise. Metabolon data analysts use proprietary visualization and interpretation software to confirm the consistency of peak identification among the various samples. Library matches for each compound were checked for each sample and corrected if necessary.

*Metabolite quantification and data normalization:* Peaks were quantified using area-under-the-curve. For studies spanning multiple days, a data normalization step was performed to correct variation resulting from instrument inter-day tuning differences. Essentially, each compound was corrected in run-day blocks by registering the medians to equal one (1.00) and normalizing each data point proportionately. For studies that did not require more than one day of analysis, no normalization is necessary, other than for purposes of data visualization.

#### Nightingale Health

Profiling of circulating lipids, fatty acids, and metabolites was done by a high-throughput targeted NMR platform (Nightingale Health© (Helsinki, Finland)) at the University of Bristol. 70 µL plasma and 70 µL sodium phosphate buffer (75 mM Na2HPO4, 0.08% sodium 3-(trimethylsilyl)propionate-2,2,3,3-d4, 0.04% sodium azide in 80%/20% H20/D2O, pH 7.4) were mixed and transferred to 3 mm NMR tubes using an 8-channel, Varispan Janus liquid handling robot (PerkinElmer).

The NMR metabolite quantification was achieved through measurements of three molecular windows from each serum/plasma sample. Two of the spectra (lipoprotein lipids (LIPO) and low molecular-weight metabolites (LMWM) windows) are acquired from native serum/plasma and one spectrum from serum lipid/plasma extracts (lipid extracts (LIPID) window). The NMR spectra were acquired using a Bruker Avance III HD 600MHz spectrometer equipped with a nitrogen-cooled triple resonance probe (CryoProbe Prodigy TCI) equipped with SampleJet auto-sampler with cooled (6°C) sample storage. Measurements of native plasma samples and plasma lipid extracts are conducted at 37°C and 22°C, respectively.

The NMR spectra were analysed for metabolite quantification (molar concentrations) in an automated fashion. For each metabolite, a ridge regression model was applied for quantification in order to overcome the problems of heavily overlapping spectral data. In the case of the lipoprotein lipid data, quantification models were calibrated using high-performance liquid chromatography methods, and individually cross-validated against NMR-independent lipid data. Low-molecular-weight metabolites, as well as lipid extract measures, were quantified as mmol/L based on regression modelling calibrated against a set of manually fitted metabolite measures. The calibration data were quantified based on iterative line-shape fitting analysis using PERCH

NMR software (PERCH Solutions Ltd., Kuopio, Finland). Quantification could not be directly established for the lipid extract measures due to experimental variation in the lipid extraction protocol. Therefore, plasma lipid extracts were scaled to total a standard serum cholesterol sample from the LIPO spectrum.

QC reports from Nightingale provide comparisons of metabolite distributions with external sources and flag potential outliers. Further description of the methodology has been published previously (Soininen *et al.*, 2009, 2015; Shin *et al.*, 2014).

#### Born in Bradford

##### Description

The BiB study is a population-based prospective birth cohort. In total, 12,453 women who experienced 13,776 pregnancies were recruited during an oral glucose tolerance test (OGTT) at approximately 26–28 weeks' gestation, which was offered to all women booked for delivery at Bradford Royal Infirmary (BRI) (with the exception of those with pre-existing diabetes (N = 70 - 0.5% of BiB pregnancies)). Eligible women had an expected delivery between March 2007 and December 2010. The study is unique because it includes high proportions of White European and South Asian families, all residing in Bradford, UK. Bradford is a city in the North of England with high levels of socioeconomic deprivation, and the cohort was started due to a high prevalence of poor child health in the city. The study website provides more information, including protocols, questionnaires and information on how researchers can access data and a full list of all available data (https://borninbradford.nhs.uk/research/). Mothers and their partners, who were recruited into the study, provided detailed interview questionnaire data, measurements, and biological samples. They also consented to the linkage of their and their child's data to routine (primary and secondary care) health and education data.

##### Details of ethics approvals

Ethical approval for the study was granted by the Bradford National Health Service Research Ethics Committee (ref 06/Q1202/48), and all participants gave written informed consent.

#### Avon Longitudinal Study of Parents and Children

##### Description of study numbers

Pregnant women residing in Avon, UK with expected dates of delivery 1st April 1991 to 31st December 1992 were invited to take part in the study. The initial number of pregnancies enrolled is 14,541 (for these at least one questionnaire has been returned or a "Children in Focus" clinic had been attended by 19/07/99). Of these initial pregnancies, there was a total of

14,676 fetuses, resulting in 14,062 live births and 13,988 children who were alive at 1 year of age.

When the oldest children were approximately 7 years of age, an attempt was made to bolster the initial sample with eligible cases who had failed to join the study originally. As a result, when considering variables collected from the age of seven onwards (and potentially abstracted from obstetric notes) there are data available for more than the 14,541 pregnancies mentioned above. The number of **new pregnancies** not in the initial sample (known as Phase I enrolment) that are currently represented on the built files and reflecting enrolment status at the age of 24 is 913 (456, 262 and 195 recruited during Phases II, III and IV respectively), resulting in an additional 913 children being enrolled. The phases of enrolment are described in more detail in the cohort profile paper and its update (see footnote 4 below). The total sample size for analyses using any data collected after the age of seven is therefore 15,454 pregnancies, resulting in 15,589 fetuses. Of these 14,901 were **alive at 1 year of age**.

A 10% sample of the ALSPAC cohort, known as the **Children in Focus (CiF) group**, attended clinics at the University of Bristol at various time intervals between 4 to 61 months of age. The CiF group were chosen at random from the last 6 months of ALSPAC births (1432 families attended at least one clinic). Excluded were those mothers who had moved out of the area or were lost to follow-up, and those partaking in another study of infant development in Avon.

Please note that the study website (http://www.bristol.ac.uk/alspac/researchers/our-data/) contains details of all the data that is available through a fully searchable data dictionary and variable search tool.

#### Data management

Study data were collected and managed using REDCap electronic data capture tools hosted at University of Bristol (Harris *et al.*, 2009). REDCap (Research Electronic Data Capture) is a secure, web-based application designed to support data capture for research studies, providing 1) an intuitive interface for validated data entry; 2) audit trails for tracking data manipulation and export procedures; 3) automated export procedures for seamless data downloads to common statistical packages; and 4) procedures for importing data from external sources.

#### Details of ethics approvals

Consent for biological samples has been collected in accordance with the Human Tissue Act (2004). Informed consent for the use of data collected via questionnaires and clinics was obtained from participants following the recommendations of the ALSPAC Ethics and Law Committee at the time. Details of ethics approvals relevant to this study: National Research Ethics Service Committee South West – Frenchay: 14/SW/1173 ALSPAC Focus at 24+ (24th

February 2015, confirmed 20th March 2015). For more information on ALSPAC's ethical procedures see: http://www.bristol.ac.uk/alspac/researchers/research-ethics/.

### Supplementary Discussion

#### Sources and properties of missingness

Variable (including very high) rates of missingness are a feature of metabolomics data relevant to quality control and as such is an important consideration in the *metaboprep* workflow. There are many reasons why metabolite measures can be missing in metabolomics datasets. Perhaps the most fundamental is that the molecules are truly absent from the sample. However, there are a range of technical reasons for missing values including instrument sensitivity thresholds that mean very low or high concentrations of particular metabolites may not be detected (typically referred to as below or above the limit of detection (LOD)) and limitations in computational processing of spectra. For a detailed discussion of this topic see (Do *et al.*, 2018).

Given the range of different reasons behind missing values, we can also expect the properties of these values to differ. Some technical issues may result in missing values that can be classed as missing completely at random (MCAR), that is, they originate from random errors during the data acquisition process. However, in many cases missing metabolite data will be either missing not at random (MNAR) such that the likelihood of a value being missing is related to the value itself (e.g., missing values caused by concentrations below the LOD) or missing at random (MAR) such that the likelihood of missingness is determined by some (observed) variable (but not related to the value itself). The classification of missing data in this way is highly relevant when considering the most appropriate way to deal with missing data, including imputation approaches (Wei *et al.*, 2018).

Variable rates of missingness (even when not excessively high) can have implications for downstream analyses. Multivariable linear models are commonly used and whilst they can run with missing data, doing so will effectively mean there is a different sample size for each metabolite.  Other approaches, such as linear mixed models and PCA, require 'complete' data to run (although adaptations such as probabilistic PCA do exist). For these reasons, many researchers choose to perform a missing value imputation step prior to their main analyses. The range and performance of imputation approaches suitable for use in metabolomics has been evaluated elsewhere (Wei *et al.*, 2018; Do *et al.*, 2018). Yet, in general, imputation can be expected to perform the best when missingness rates are relatively low and the correlation between variables relatively high. For this reason, whilst it clearly brings benefit in terms of delivering a 'complete' dataset, the actual gain in information content within the context of metabolomics can be expected to be small. Importantly, more robust methods for imputation, such as multivariable multiple imputation, can only be undertaken during subsequent main analyses (addressing a specific question), as these require data included in main analysis models to be included in the imputation process (Hughes *et al.*, 2019).

### Supplementary Figures

#### Figure S1. Born in Bradford sample selection flow chart

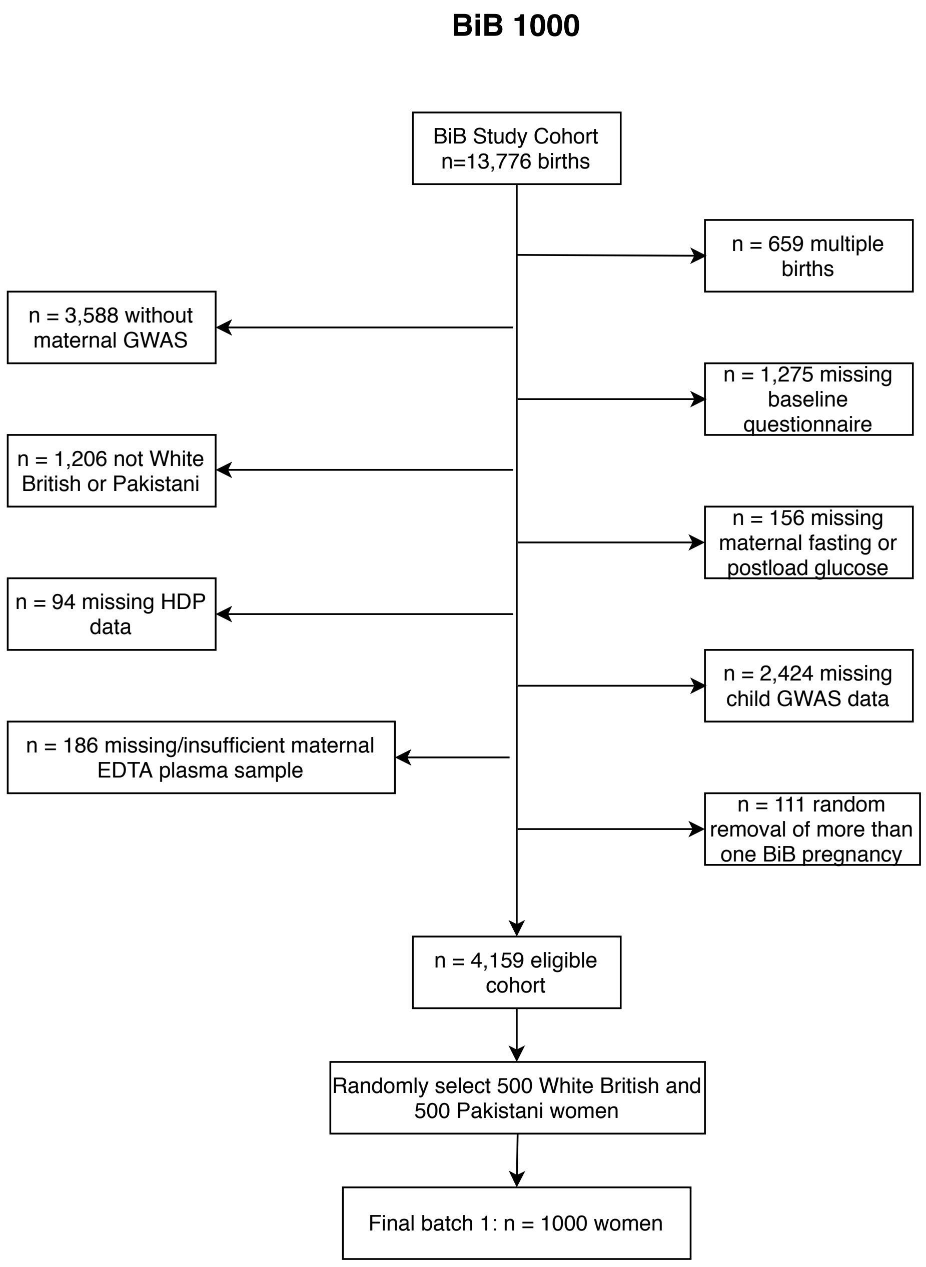

#### Supplementary Tables

##### Table S1. Born in Bradford (BiB) dataset: Participant characteristics

| | | Batch 1 (N = 1000) | Batch 2 (N= 2000) |
|---|---|---|---|
| **Maternal Characteristics** | **Category** | - | - |
| ***Age, years*** | | 27.5 (5.7) | 27.5 (5.7) |
| ***Parity*** | Nulliparous | 359 (35.9) | 745 (37.3) |
| | Multiparous | 611 (61.1) | 1213 (60.7) |
| Missing (%) | | 30 (3.0) | 42 (2.1) |
| ***BMI (kg/m²)*** | | 26.7 (6.0) | 26.8 (5.9) |
| Missing (%) | | 36 (3.6) | 97 (4.9) |
| ***Ethnicity*** | White British | 500 (50.0) | 933 (46.7) |
| | Pakistani | 500 (50.0) | 1067 (53.3) |
| ***HDP*** | Normotensive | 885 (88.5) | 1690 (84.5) |
| | PE | 25 (2.5) | 79 (4.0) |
| | GHT | 69 (6.9) | 231 (11.6) |
| Missing (%) | | 21 (2.1) | - |
| ***Gestational Diabetes*** | Yes | 91 (9.1) | 264 (13.2) |
| ***IMD*** | Quintile 1 | 656 (65.6) | 1340 (67.0) |
| | Quintile 5 | 19 (1.9) | 37 (1.8) |
| ***Smoking*** | Yes | 176 (17.6) | 378 (18.9) |
| Missing (%) | | 1 (0.1) | 3 (0.2) |
| ***Alcohol*** | Yes | 338 (33.8) | 630 (31.5) |
| Missing (%) | | 1 (0.1) | 3 (0.2) |
| ***Gest Age at Blood Sampling (weeks)*** | | 26.2 (2.0) | 26.2 (2.0) |

Data are means ± SD or n (%) unless stated.
For characteristics with no “missing” category, data were 100% complete.

#### Table S2. Avon Longitudinal Study of Parents and Children (ALSPAC) dataset: Participant characteristics

| | |
|---|---|
| No. of samples | 3361 |
| No. of unique individuals[a] | 3277 |
| Sex, female (%) | 1973 (60.2%) |
| Ethnicity[b], non-white (%) | 108 (3.7%) |
| Missing (%) | 371 (11.3%) |
| Age[c], years (mean, SD) | 24.0 (0.8) |
| Missing (%) | 1 (0.03%) |
| Body mass index[c], kg/m$^2$ (mean,SD) | 24.7 (4.9) |
| Missing (%) | 34 (1.0%) |
| Smoking[c], daily smokers (%) | 381 (11.8%) |
| Missing (%) | 52 (1.6%) |

[a] replicate samples were analysed for 84 individuals (all summary statistics are based on unique individuals); [b] ethnicity (white/non-white) as defined based on mother's self-reported ethnicity and that of her partner (ascertained during gestation); [c] based on data collected during the same clinic as samples were collected (F24). SD = standard deviation.
