## Supplementary_Data_2 for "*metaboprep*: an R package for pre-analysis data description and processing"

### metaboprep metabolomics data preparation report

2021-07-05

metaboprep report relates to:

- Project: BiB_MS-1
- Platform: Metabolon

The `metaboprep R` package performs three operations:

1. Provides an assessment and summary statistics of the raw metabolomics data.
2. Performs data filtering on the metabolomics data.
3. Provides an assessment and summary statistics of the filtered metabolomics data, particularly in the context of batch variables when available.

This report provides descriptive information for raw and filtered metabolomics data for the project BiB_MS-1.

The data filtering workflow is as follows:

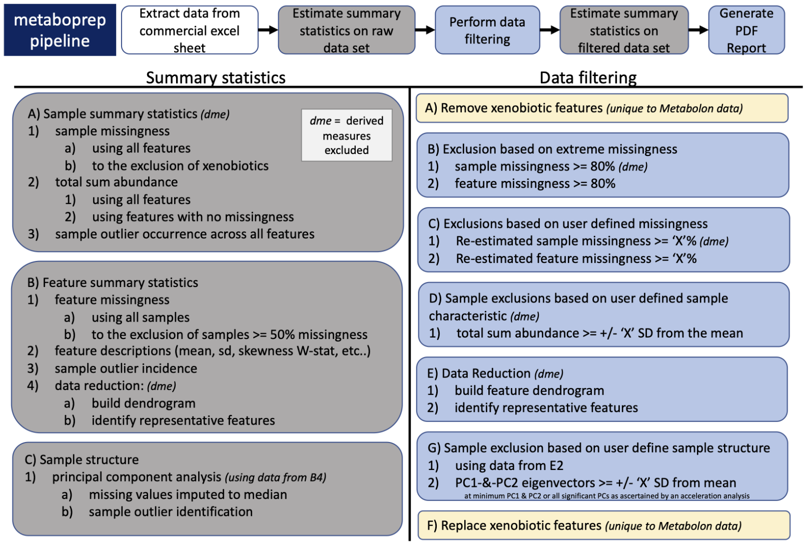


1. Issues can be raised on GitHub.
2. Questions relating to the `metaboprep` pipeline can be directed to David Hughes:.
3. `metaboprep` is published in Journal to be determined and can be cited as:

### 1 Sample size of BiB_MS-1 data set

| data.set | raw.data | filtered.data |
|---|---|---|
| number of samples | 1000 | 989 |
| number of features | 1369 | 1037 |

#### 1.1 Missingness

Missingness is evaluated across samples and features using the original/raw data set.

##### 1.1.1 Visual structure of missingness in your raw data set.

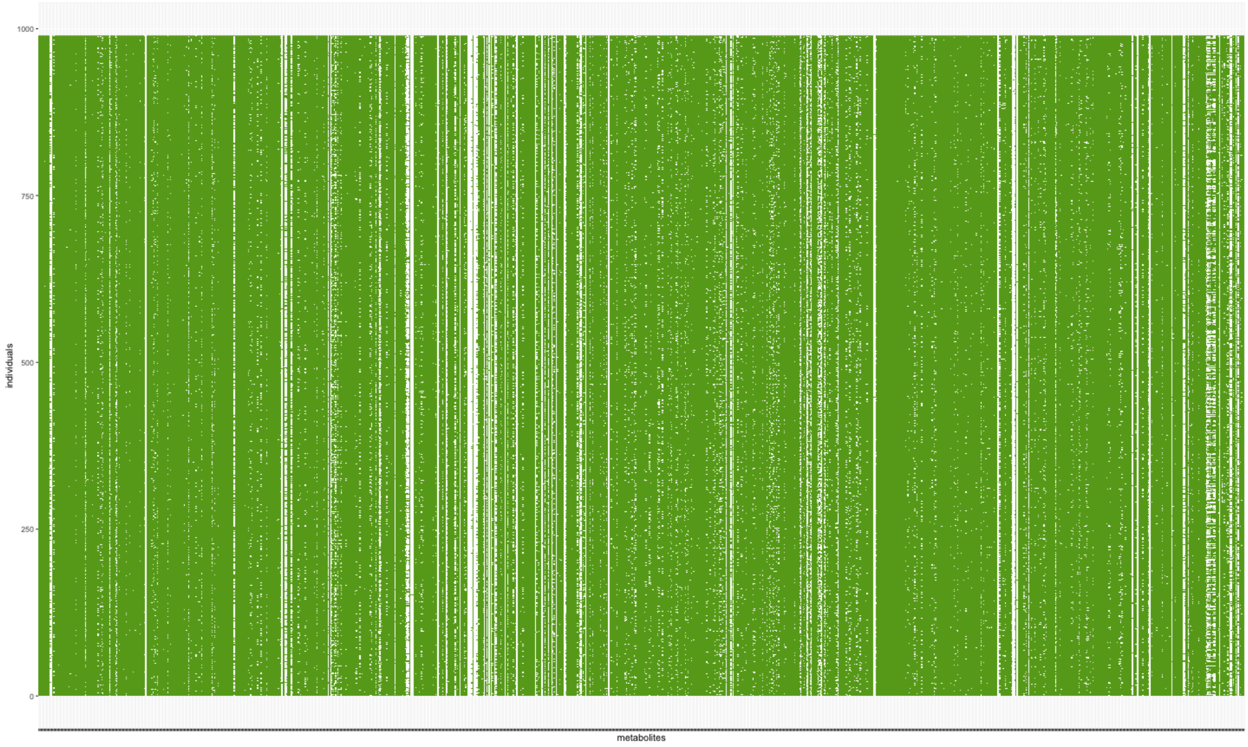


**Figure Legend:** Missingness structure across the raw data table. White cells depict missing data. Individuals are in rows, metabolites are in columns.

##### 1.1.2 Summary of sample and feature missingness

-- Initial raw data set: Estimates of missingness for samples and features --

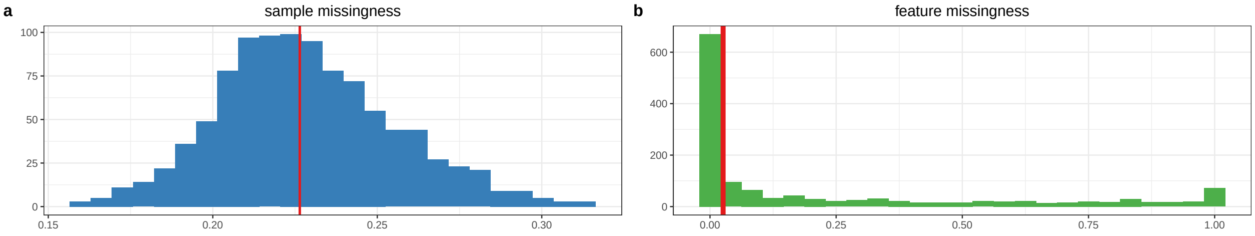


c

| percentile | samples | features |
|---|---|---|
| 0 | 0.1578 | 0 |
| 0.25 | 0.2104 | 0 |
| 0.5 | 0.2264 | 0.026 |
| 0.75 | 0.2454 | 0.381 |
| 1 | 0.3112 | 1 |

d

| percent.missing | sample.size |
|---|---|
| 10% feature missingness | 900 |
| 20% feature missingness | 800 |
| 30% feature missingness | 700 |
| 40% feature missingness | 600 |
| 50% feature missingness | 500 |

**Figure Legend:** Raw data - (a) Distribution of sample missingness with sample mean illustrated by the red vertical line. (b) Distribution of feature missingness sample mean illustrated by the red vertical line. (c) Table of sample and feature missingness percentiles. A tabled version of plot a and b. (d) Estimates of study samples sizes under various levels of feature missingness.

#### 1.2 Data Filtering

##### 1.2.1 Exclusion summary

| exclusions | count |
|---|---|
| Extreme_sample_missingness | 0 |
| Extreme_feature_missingness | 96 |
| User_defined_sample_missingness | 0 |
| User_defined_feature_missingness | 236 |
| User_defined_sample_totalpeakarea | 0 |
| User_defined_sample_PCA_outliers | 11 |

**Table Legend**: Six primary data filtering exclusion steps were made during the preparation of the data. (1) Samples with missingness >=80% were excluded. (2) features with missingness >=80% were excluded (xenobiotics are not included in this step). (3) sample exclusions based on the user defined threshold were excluded. (4) feature exclusions based on user defined threshold were excluded (xenobiotics are not included in this step). (5) samples with a total-peak-area or total-sum-abundance that is >= N standard deviations from the mean, where N was defined by the user, were excluded. (6) samples that are >= N standard deviations from the mean on principal component axis 1 and 2, where N was defined by the user, were excluded.

##### 1.2.2 Metabolite or feature reduction and principal components

A data reduction was carried out to identify a list of representative features for generating a sample principal component analysis. This step reduces the level of inter-correlation in the data to ensure that the principal components are not driven by groups of correlated features.

| data.reduction | count |
| --- | --- |
| total metabolite count | 858 |
| metabolites included in data reduction | 858 |
| number of metabolite clusters | 483 |
| number of representative metabolites | 483 |

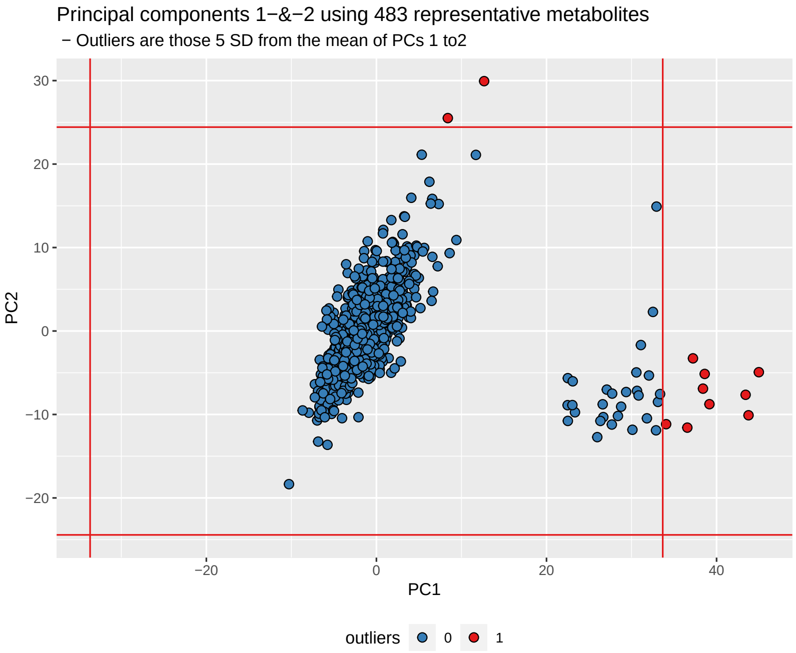


**Figure Legend:** The data reduction table on the left presents the number of metabolites at each phase of the data reduction (Spearman's correlation distance tree cutting) analysis. On the right principal components 1 and 2 are plotted for all individuals, using the representitve features identified in the data reduction analysis. The red vertical and horizontal lines indicate the standard deviation (SD) cutoffs for identifying individual outliers, which are plotted in red. The standard deviations cuttoff were defined by the user.

---

### 2 Filtered data

## 2.1 N

- The number of samples in data = 989
- The number of features in data = 1037

#### 2.2 Relative to the raw data

- 11 samples were filtered out, given the user's criteria.
- 332 features were filtered out, given the user's criteria.
- Please review details above and your log file for the number of features and samples excluded and why.

#### 2.3 Summary of filtered data

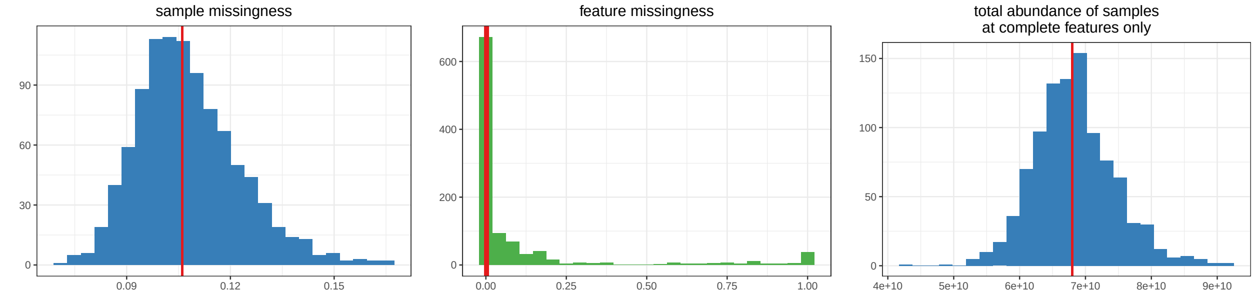


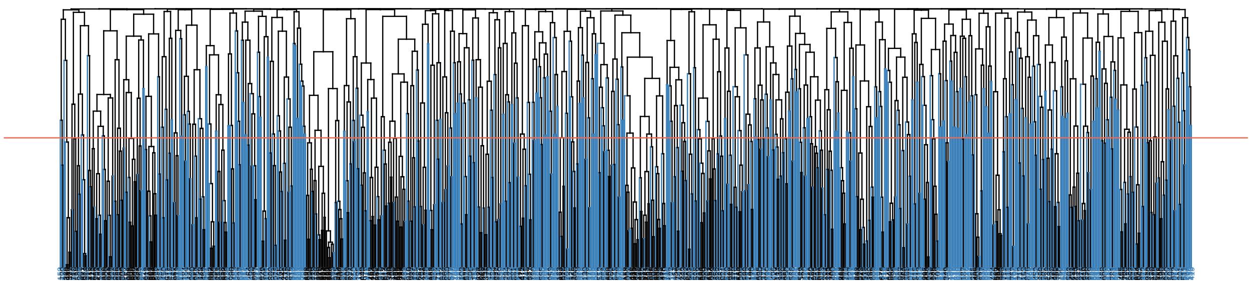

| data.reduction | count |
|---|---|
| total metabolite count | 1037 |
| metabolites included in data reduction | 919 |
| number of metabolite clusters | 512 |
| number of representative metabolites | 511 |

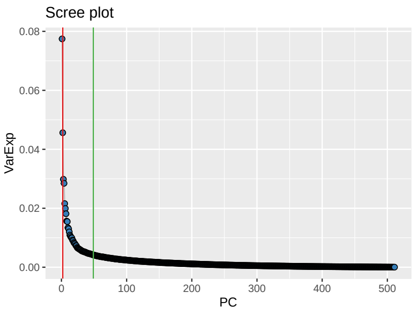


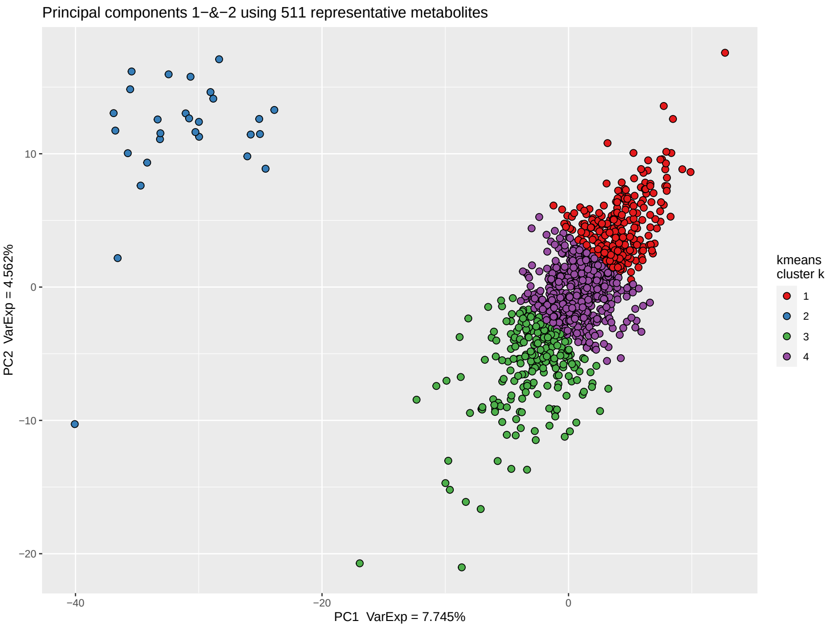


**Figure Legend:** Filtered data summary. Distributions for sample missingness, feature missingness, and total abundance of samples. Row two of the figure provides a Spearman's correlation distance clustering dendrogram highlighting the metabolites used as representative features in blue, the clustering tree cut height is denoted by the horizontal line. Row three provides a summary of the metabolite data reduction in the table, a Scree plot of the variance explained by each PC and a plot of principal component 1 and 2, as derived from the representative metabolites. The Scree plot also identifies the number of PCs estimated to be informative (vertical lines) by the Cattel's Scree Test acceleration factor (red, n = 2) and Parallel Analysis (green, n = 49). Individuals in the PC plot were clustered into 4 kmeans (k) clusters, using data from PC1 and PC2. The kmeans clustering and color coding is strictly there to help provide some visualization of the major axes of variation in the sample population(s).

#### 2.4 Structure among samples: top 5 PCs

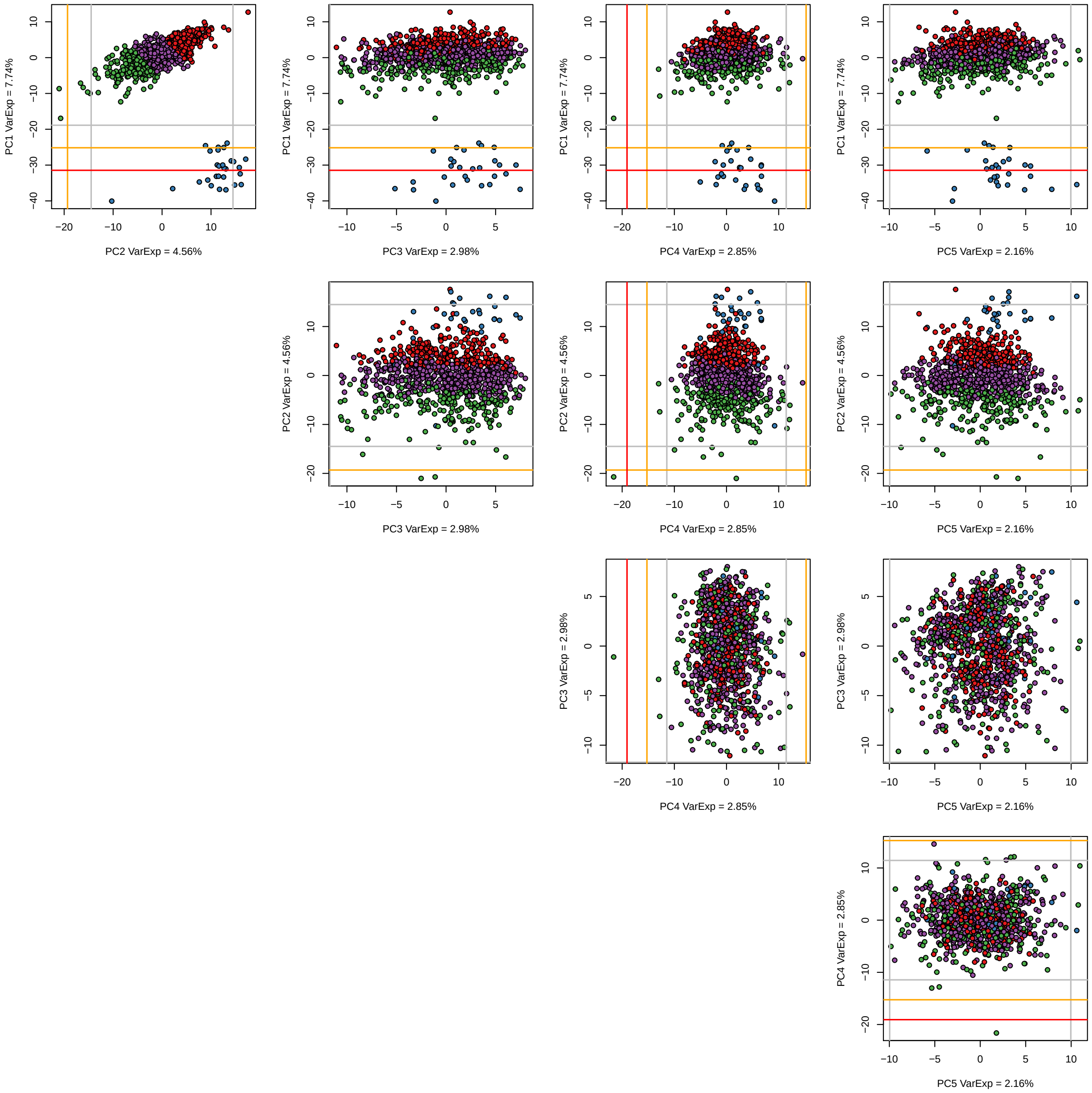


**Figure Legend:** A matrix plot of the top five principal components including demarcations of the 3rd (grey), 4th (orange), and 5th (red) standard deviations from the mean. Samples are color coded as in the summary PC plot above using a kmeans analysis of PC1 and PC2 with a k (number of clusters) set at 4. The choice of k = 4 was not robustly chosen it was a choice of simplicity to help aid visualize variation and sample mobility across the PCs.

#### 2.5 Feature Distributions

##### 2.5.1 Estimates of normality: W-statistics for raw and log transformed data

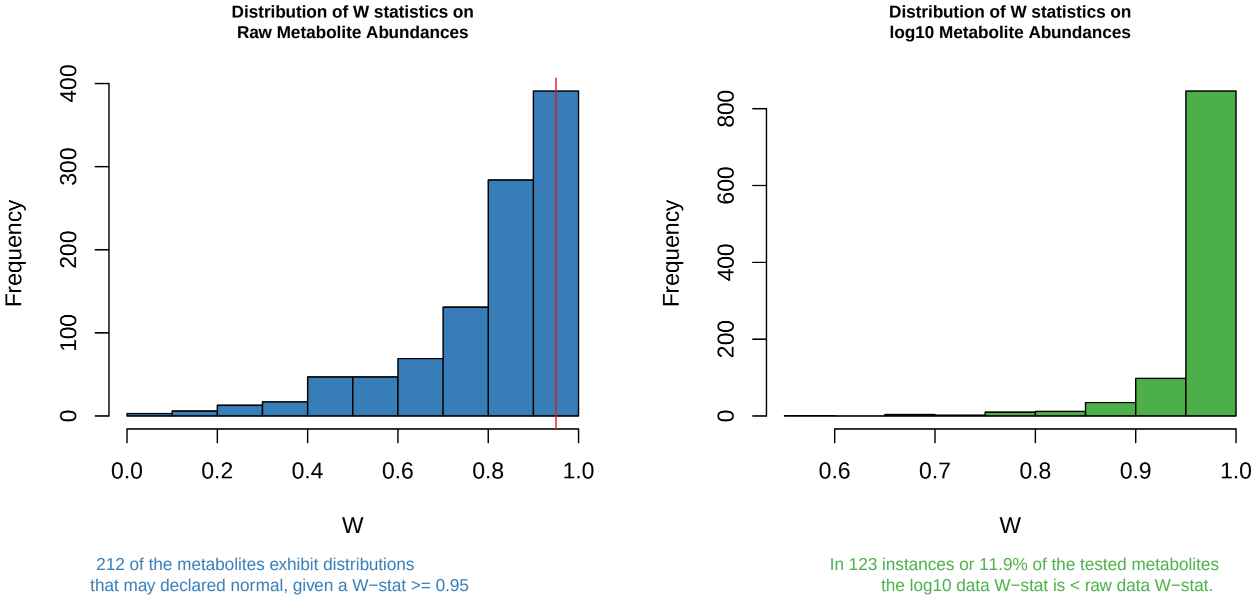


**Figure Legend:** Histogram plots of Shapiro W-statistics for raw (left) and log transformed (right) data distributions. A W-statistic value of 1 indicates the sample distribution is perfectly normal and value of 0 indicates it is perfectly uniform. Please note that log transformation of the data *may not* improve the normality of your data.

**Analysis details:** Of the 1037 features in the data 29 features were excluded from this analysis because of no variation or too few observations ($n < 40$). Of the remaining 1008 metabolite features, a total of 212 may be considered normally distributed given a Shapiro W-statistic >= 0.95.

##### 2.5.2 Distributions

A pdf report is being written to BiB_MS-1_outlier_detection_pre_filtering.pdf that contains dotplot, histogram and distribution summary statistics for each metabolite in your data set, providing an opportunity to visually inspect all your metabolites feature data distributions.

#### 2.6 Outliers

**Evaluation of the number of samples and features that are outliers across the data.**

| percentile | outlying.features.by.sample | outlying.samples.by.feature |
|---|---|---|
| 0% | 0 | 0 |
| 25% | 0 | 1 |
| 50% | 1 | 2 |
| 75% | 3 | 4 |
| 100% | 44 | 15 |

**Table Legend:** The table reports the number of point estimates for the minimum (0%) median (50%) and maximum (100%) number of outlying features across samples and the number of outlying samples across features.

##### 2.6.1 Notes on outlying samples at each metabolite|feature

There may be extreme outlying observations at individual metabolites|features that have not been accounted for. You may want to:

1. Turn these observations into NAs.
2. Winsorize the data to some maximum value.
3. Rank normalize the data which will place those outliers into the top of the ranked standard normal distribution.
4. Turn these observations into NAs and then impute them along with other missing data in your data set.

---

### 3 Influence of batch variables on filtered data

#### 3.1 Filtered data *feature* missingness: influenced by possible explanatory variables

Feature missingness may be influenced by the metabolites (or features) biology or pathway classification, or your technologies methodology. The figure(s) below provides an illustrative evaluation of the proportion of *feature missigness* as a product of the variable(s) available in the raw data files.

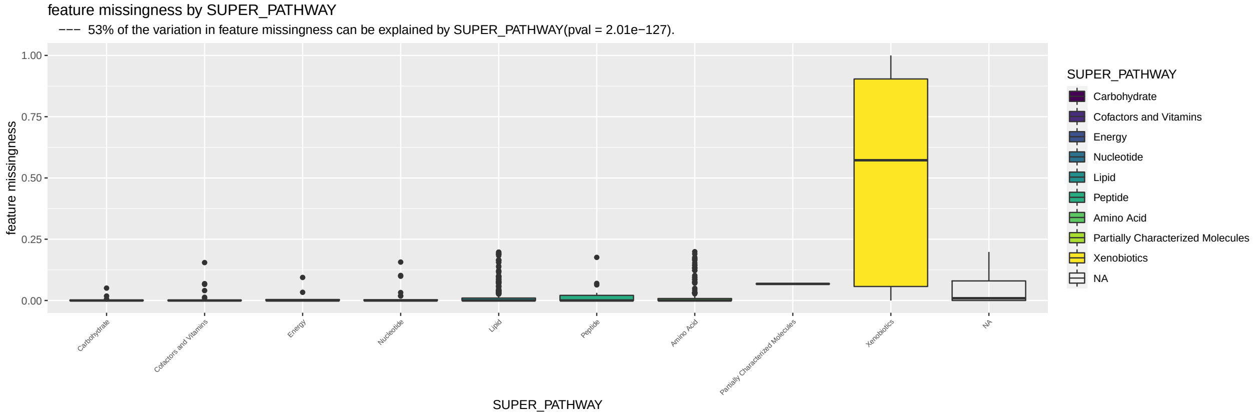


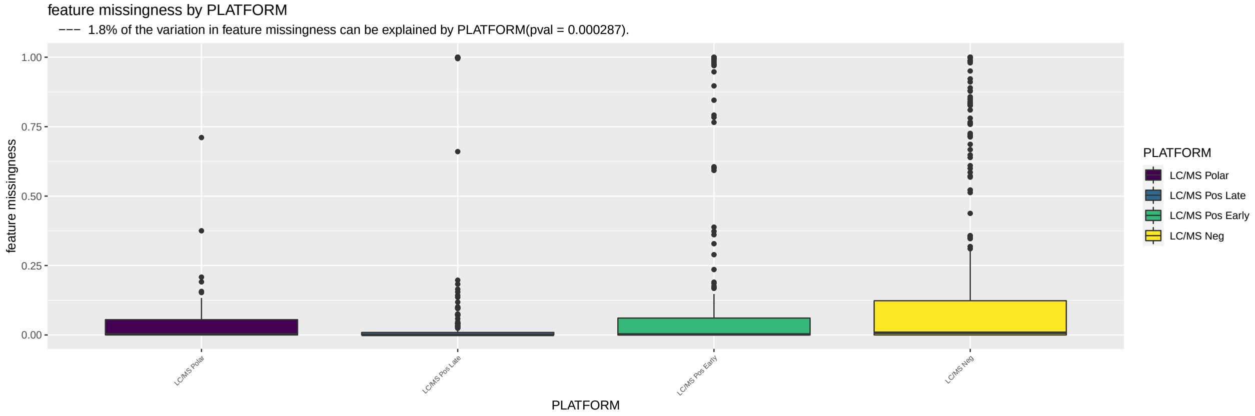


**Figure Legend:** Box plot illustration(s) of the relationship that available batch and biological variables have with feature missingness.

#### 3.2 Filtered data *sample* missingness: influenced by possible explanatory variables

The figure provides an illustrative evaluation of the proportion of *sample missingness* as a product of sample batch variables provided by your supplier. This is the univariate influence of batch effects on *sample missingness*.

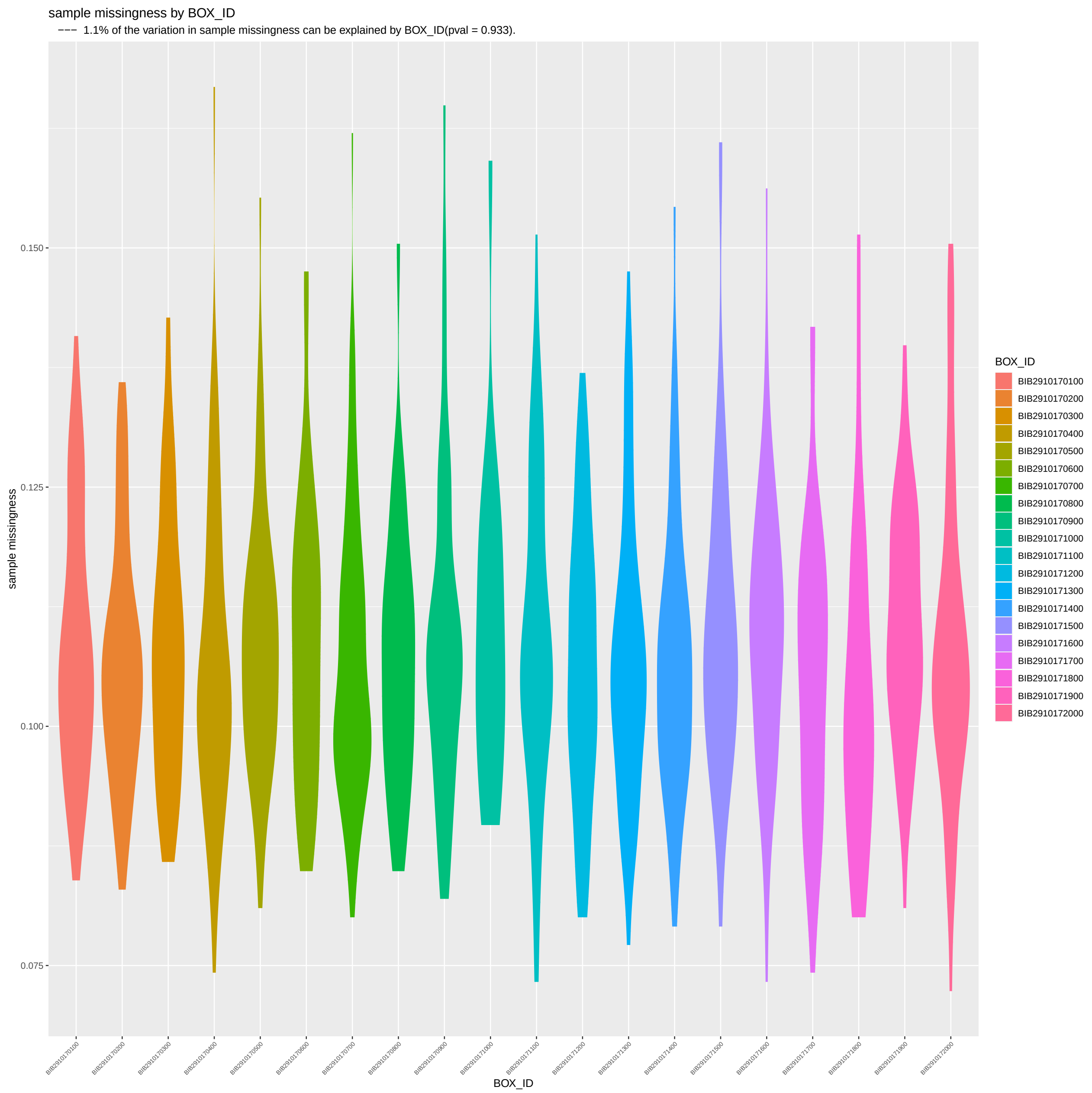


**Figure Legend:** Box plot illustration(s) of the relationship that available batch variables have with sample missingness.

#### 3.3 Multivariate evaluation: batch variables

| batch.variable | etasq.var.exp | pvalue |
|---|---|---|
| box.id | 1.09 | 9.33e−01 |
| residuals | 98.91 | NA |

**Table Legend:** TypeII ANOVA: the eta-squared (eta-sq) estimates are an estimation of the percent of variation explained by each independent variable, after accounting for all other variables, as derived from the sum of squares. This is a multivariate evaluation of batch variables on *sample missingness*.

### 4 Sample Total Peak|Abundance Area (TPA):

Total peak|abundance area (TPA) is simply the sum of the abundances measured across all features. TPA is one measure that can be used to identify unusual samples given their entire profile. However, the level of missingness in a sample may influence TPA. To account for this we:

1. Evaluate the correlation between TPA estimates across all features with TPA measured using only those features with complete data (no missingness).

2. Determine if the batch effects have a measurable impact on TPA.

#### 4.1 Relationship with missingness

Correlation between total peak area (at complete features) and missingness

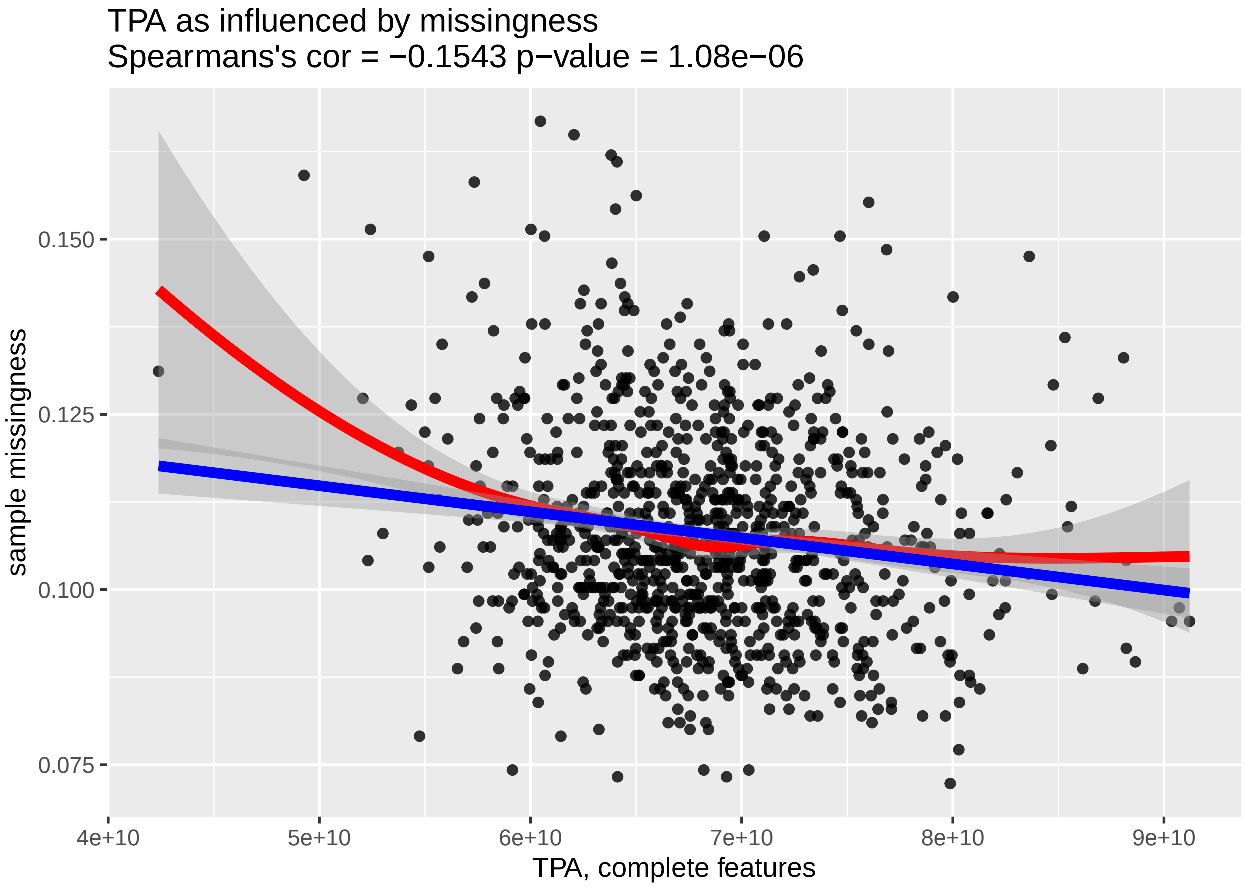


**Figure Legend:** Relationship between total peak area at complete features (x-axis) and sample missingness (y-axis).

#### 4.2 Univariate evaluation: batch effects

The figure below provides an illustrative evaluation of the *total peak area* as a product of sample batch variables provided by your supplier.

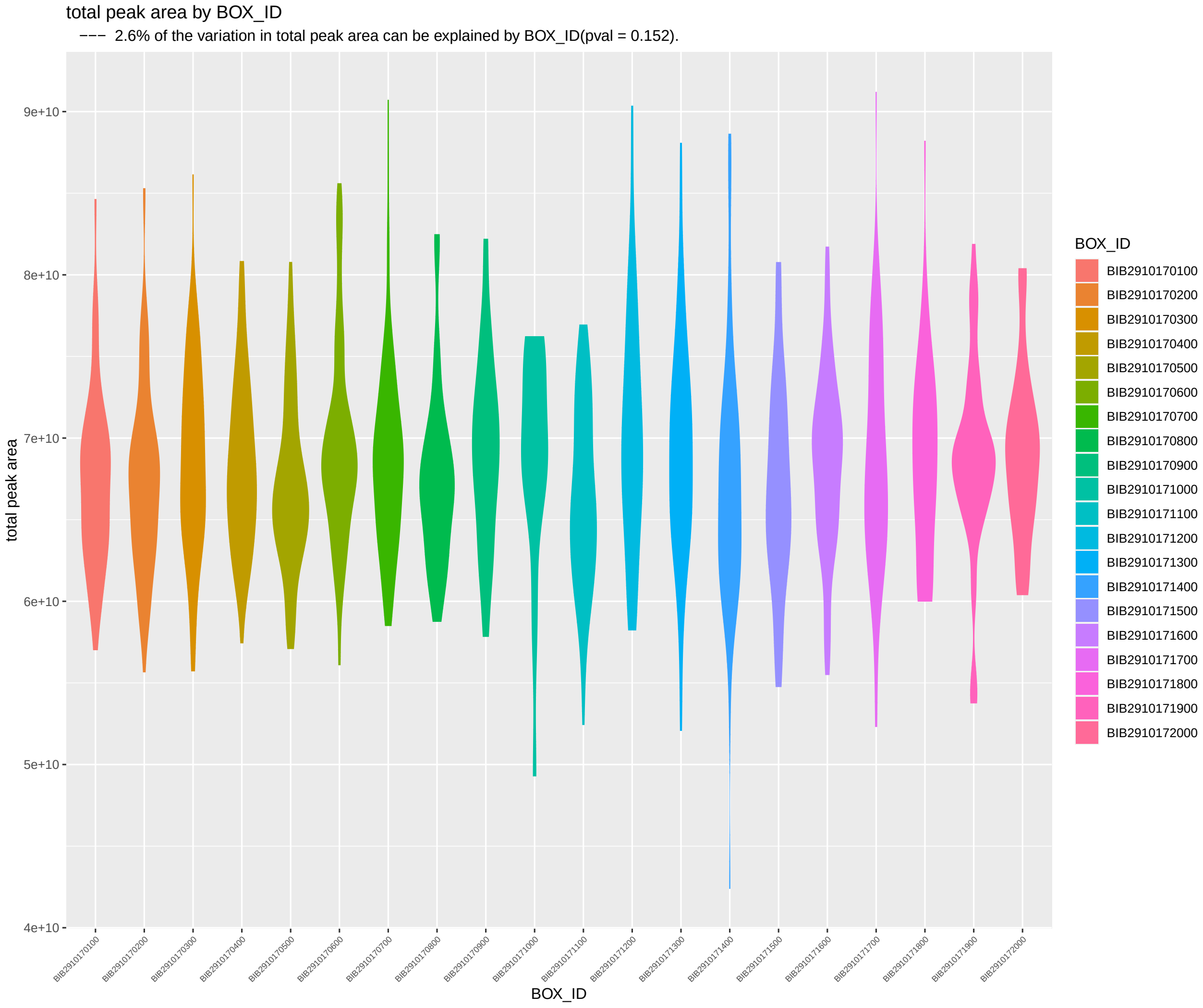


**Figure Legend:** Violin plot illustration(s) of the relationship between total peak area (TPA) and sample batch variables that are available in your data.

##### 4.2.1 Multivariate evaluation: batch variables

| batch.variable | etasq.var.exp | pvalue |
|---|---|---|
| box.id | 2.55 | 1.52e−01 |
| residuals | 97.45 | NA |

**Table Legend:** TypeII ANOVA: the eta-squared (eta-sq) estimates are an estimation on the percent of variation explained by each independent variable, after accounting for all other variables, as derived from the sum of squares. This is a multivariate evaluation of batch variables on *total peak|abundance area* at complete features.

### 5 Power analysis

##### Exploration for case/control and continuous outcome data using the filtered data set

Analytical power analysis for both continuous and imbalanced presence/absence correlation analysis.

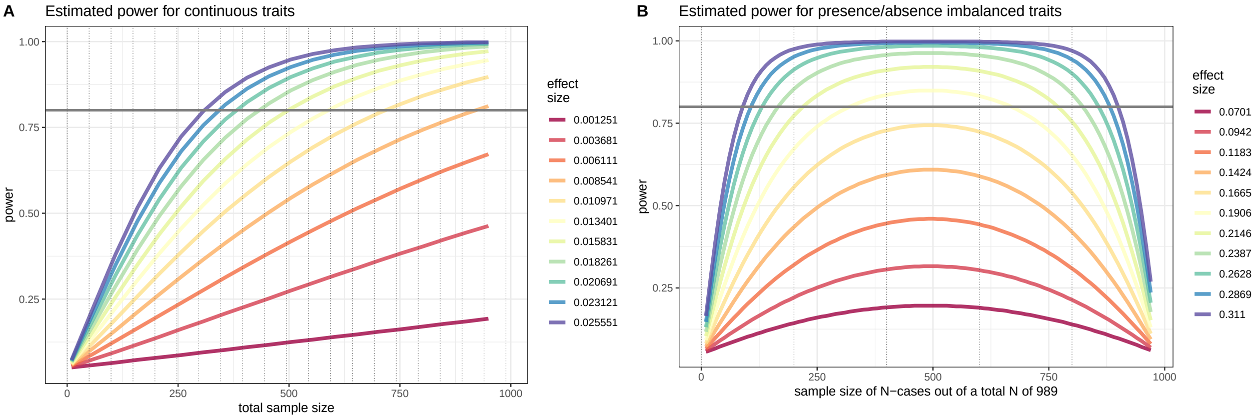


**Figure Legend:** Simulated effect sizes are illustrated by their color in each figure. Figure (A) provides estimates of power for continuous traits with the total sample size on the x-axis and the estimated power on the y-axis. Figure (B) provides estimates of power for presence/absence (or binary) traits in an imbalanced design. The estimated power is on the y-axix. The total sample size is set to 989 and the x-axis depicts the number of individuals present (or absent) for the trait. The effects sizes illustrated here were chosen by running an initial set of simulations which identified effects sizes that would span a broad range of power estimates given the sample population's sample size.
